# Photon-statistics in sensitized emission FRET and FLIM: a comparative theoretical analysis

**DOI:** 10.1101/774919

**Authors:** Alessandro Esposito

## Abstract

FRET imaging is an essential analytical method in biomedical research. The limited photon-budget experimentally available, however, imposes compromises between spatiotemporal and biochemical resolutions, photodamage and phototoxicity. The study of photon-statistics in biochemical imaging is thus important in guiding the efficient design of instrumentation and assays. Here, we show a comparative analysis of photon-statistics in FRET imaging demonstrating how the precision of FRET imaging varies vastly with imaging parameters. Therefore, we provide analytical and numerical tools for assay optimization. FLIM is a very robust technique with excellent photon-efficiencies but also intensity-based FRET imaging can reach very high precision by utilizing also information within acceptor fluorescence.

## 1. Introduction

Förster resonance energy transfer (FRET) is the non-radiative transfer of energy from a donor fluorophore to an acceptor chromophore [1, 2]. The probability for a molecule to transfer energy via FRET (*E*, FRET efficiency) depends on the inverse of the sixth power of the inter-chromophore distance and, typically, FRET efficiency is sensitive to distances within the nanometers range [3]. For its high sensitivity at the nanometer scale, FRET has many applications in biophysics and biomedical sciences, for instance, to determine intermolecular distances and protein conformational changes, to detect protein-protein interactions, protein modifications and to engineer biosensors for the detection of biomolecules (reviewed in [4–7]). FRET results in the reduction of the quantum yield and the fluorescence lifetime of the donor fluorophore; in the instances where the acceptor is a fluorescent molecule, FRET also causes the sensitized emission of fluorescence from the acceptor fluorophore [1]. Fluorescence lifetime imaging microscopy (FLIM) [8, 9] is thus one of the methodologies that enables researcher to quantitate FRET; among the various types of FLIM techniques, time-correlated single-photon counting (TCSPC) is considered the gold standard for its high precision and accuracy [10, 11]. Biological applications of FRET and FLIM are often constrained by the limited available photon-budget, *i.e.* the number of photons that can be detected within a reasonable exposure time limited by the need to avoid photodamage and phototoxicity or by the time resolution required to characterize dynamic biological processes. The role of photon-statistics for several FRET imaging techniques has been characterized, more extensively for FLIM applications [8, 12–19] and, to our knowledge, at a lesser extent for intensity-based FRET imaging techniques [20, 21].

A method that is commonly used to quantify FRET efficiencies by sensitized emission FRET (seFRET) relies on the acquisition of three images [22, 23]: i) the image of the donor excited at a wavelength optimized for donor excitation (I^DD^), ii) the image of the acceptor acquired exciting the sample at the same wavelength used for donor excitation (I^DA^) and iii) the image of the acceptor excited at a wavelength optimized for the excitation of the acceptor (I^AA^). Usually, the selective excitation of a donor fluorophore (avoiding excitation of the acceptor fluorophore) or the selective detection of the FRET-sensitized emission of the acceptor fluorophore (with no direct excitation or no contamination from the fluorescence emitted by the donor fluorophore) is not possible. Therefore, several groups have developed methods and software for the correction of spectral cross-talks [22–26] or spectral unmixing [27–30]. For example, we can estimate the donor spectral bleed-through into the acceptor channel using a donor-only control sample and measuring the ratio of intensities detected in the acceptor an donor channels: DER = [I^DA^/I^DD^]_only-donor_ (donor emission ratio). Similarly, we can estimate the direct excitation of acceptor fluorophores by exciting an acceptor-only sample with excitation light optimal for acceptor excitation, and measuring the ratio of intensities detected into the two channels: AER = [I^DA^/I^AA^]_only-acceptor_ (acceptor excitation ratio). The corrected FRET signal (cFRET) can be then computed on each pixel of the sample where both donor and acceptor fluorophores are present [22, 23]:

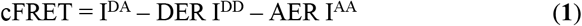

We can quantitate FRET efficiency by normalizing the FRET-sensitized fluorescence intensity cFRET to the intensity that would have been emitted by the donor if FRET did not occur (dFRET estimator) or to the intensity that would have been emitted by the acceptor if FRET efficiency was equal to 100% (aFRET estimator):

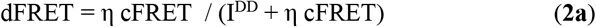

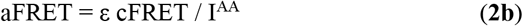

ε and η are two additional correction factors that depend on the imaging parameters and can be determined by imaging a sample exhibiting a known FRET efficiency (see also supplementary tables 1-3 to compare different nomenclatures). It has been shown [22] that dFRET and aFRET are good estimators for the apparent FRET efficiency, *i.e.*, the FRET efficiency (E) multiplied by the fraction of interacting donors (f_D_) or interacting acceptors (f_A_), respectively. Protocols for seFRET estimation are described elsewhere [22, 23, 31]. Here, we study the role of photon-statistics in seFRET and provide a theoretical comparison of the physical limits in precision and accuracy between seFRET and the well-characterized TCSPC. Interestingly, seFRET performs very well from a theoretical perspective, resulting in high precision because of the efficient utilization of information inferred from both donor and acceptor signals.

**Table 1.** properties of FRET pairs relevant to seFRET

| FRET pair | Microscope | AER | DER | $\eta$ | $\varepsilon$ | Reference |
| --- | --- | --- | --- | --- | --- | --- |
| CFP-YFP | Confocal (system 1) | 0.60 | 0.42 | 0.52 | 6.3 | [22] |
| CFP-citrine | Wide-field (system 2) | 0.29 | 1.07 | 0.015 | 42 | [23] |

## 2. Results

### A. Fisher information matrix and seFRET

We describe the wavelength-dependent (*λ*) fluorescence emission as the sum of photons emitted by the donor fluorophore, photons emitted by sensitised acceptors (SE) and photons emitted by acceptors upon direct excitation (DE) with donor excitation light source:

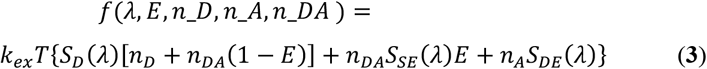

*S*_*D*_, *S*_*SE*_ and *S*_*DE*_ depend on the spectral characteristics of the fluorophores and detection system, *e.g.*, quantum yields, molar extinction coefficients, spectral overlaps; *n*_*D*_, *n*_*A*_ and *n*_*DA*_ are the number of non-interacting donor molecules, non-interacting acceptor molecules and interacting donor-acceptor pairs in the sample, respectively. After integration of Eq. 3 over the spectral bands of the donor ([λ_d1_, λ_d2_]) and the acceptor channels ([λ_a1_, λ_a2_]), it is possible to express the detected intensity in the form:

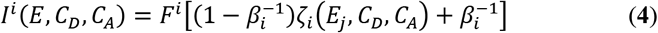

where i = DD, DA, AA and j = D, A; β_i_ is the fractional contribution of an unspecific background signal (B_i_), F^i^ is the maximum intensity of the channel relative to the value of the function ζ_i_(E,C_D_,C_A_) which is the only expression that explicitly depends on the energy transfer efficiency, the concentrations of donor (C_D_) and acceptor (C_A_) fluorophores; furthermore E_D_ = f_D_E and E_A_ = f_A_E are useful parametrization to analyze the performances of dFRET and aFRET, respectively. For both dFRET and aFRET (j = D or A), the values of F^i^, β_i_ and ζ_AA_ are (see Supp. Eq. S28-32 in supporting information):

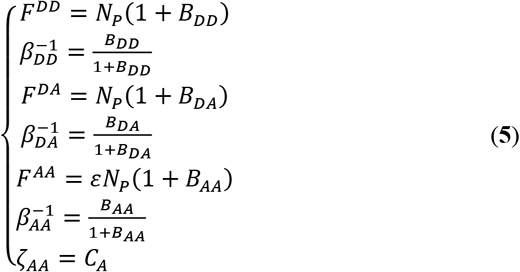

For the dFRET (Eq. 13a) and the aFRET (Eq. 13b) estimators, ζ_DD_ and ζ_DA_ are:

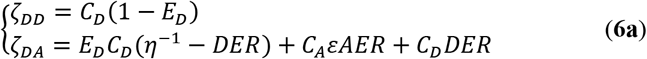

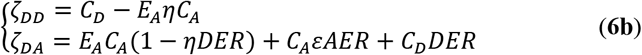

The proportionality constants ε and η (see Supp. Eq. S25-26 and Supp. Tables 1-3) are the ratio of the donor/acceptor excitation light intensities and detection efficiencies, respectively; N_P_ = k_ex_ T is the sum of photons collected in the donor and sensitized emission channel. With Eqs. 6, we can evaluate the Fisher information matrix *J* which element (J^−1^)_11_ of its inverse matrix provides the Cramer-Rao bound [20, 32] for the variance of dFRET and aFRET (see Supp. Eqs. S33-43). With the subscript B, SBT and E, we indicate the variances caused by background signals, spectral bleed-through and FRET efficiency, respectively. The variance of dFRET can be thus written as:

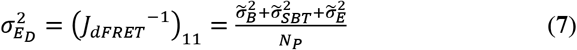

The symbol 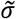 indicate variances normalized to the Poisson noise variance (N_P_ = k_ex_ T). The background, spectral bleed-through and intrinsic noise contributions are defined by Eqs. 8–10, respectively.

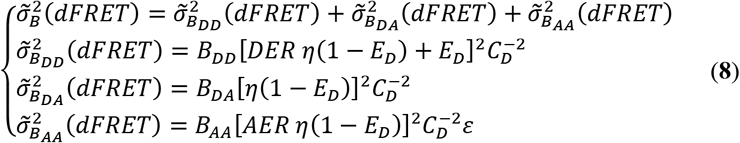

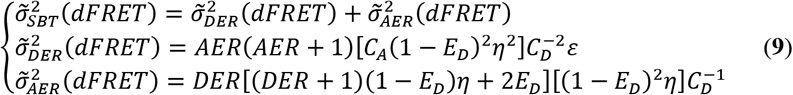

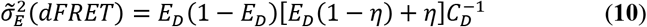

Similarly, we can evaluate the analytical descriptions for the noise of the estimator aFRET:

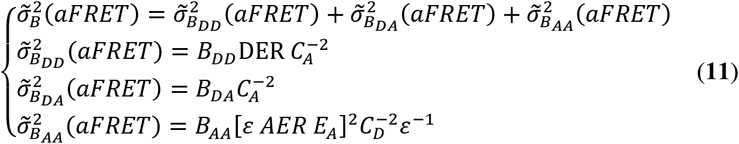

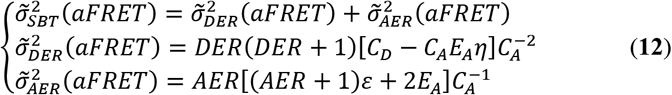

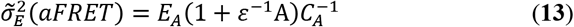

Our analytical framework is explicitly derived from the seminal work of Watkins and colleagues for sensitized emission FRET in single molecule applications [20]. In the present work, the main difference is that we study Fisher information for a typical three-filter method [22, 23, 25, 33], where we acquire the three images I^DD^, I^DA^ and I^AA^. The measurement I^AA^ was not included in the previous analysis – where was unnecessary – but it has significant implications for the photon-efficiency of seFRET. In sections C-E, these analytical solutions are tested in the absence of cross-talk, in the presence of spectral bleed-through and in the presence of unspecific channel background by means of open-source numerical simulations (see Methods).

### B. Cramer-Rao lower bound for TCSPC

To provide a reference for the theoretical efficiency of seFRET, we studied the relative error for the estimation of FRET by TCSPC, the gold-standard in FLIM detection and estimation of FRET [7]. We could not calculate the analytical solutions for an appropriate double-exponential model for TCSPC, thus we implemented open-source numerical simulations (see Methods) adapted the computational core originally developed by Bouchet and colleagues [32]. The Cramer-Rao lower bound for the standard deviation of the FRET estimate 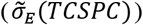, normalized to the total photon count, is shown in figure 1. We consider the case where FRET is estimated by a double exponential fit with a known reference fluorescence lifetime and where the fractional contribution of the FRET-decays and its shortened lifetime are fitted. In fig. 1a we assumed an ideal Dirac-like instrument response function (IRF), for a reference lifetime of 1, 3 and 10 nanoseconds. For each reference lifetime, we simulated a contribution of 10% (higher curves), 50% (middle curves) and ~90% (lower curves) to the total signal arising from the short exponential decay. As expected, for larger reference fluorescence lifetime values and larger relative contribution of the FRET fraction, the normalized standard deviation is lower. Fig. 1b shows the same analysis but with a finite IRF as defined in Bouchet *et al.* [32] to ~38ps full-width half-maximum. The IRF seems to not have a significant impact except for high FRET efficiencies values, *i.e.* when the fluorescence lifetime estimates are in the order of magnitude of the IRF. Fig. 1c shows similar simulation, however, in this case the reference fluorescence lifetime is kept constant to 3ns for all curves. Here, we changed the contribution of an uncorrelated background to 0 (fig. 1c, lower curves), 100 (middle curves) and 1,000 (higher curves) background photons. Within the set parameters, the intermediate signal-to-background ratio correspond to 1, 5 and 100 and the conditions with the highest background signal to 0.1, 0.5 and 1, demonostrating that the statistical error in FRET estimates are comparatively robust to the presence of uncorrelated background.

**Fig. 1.**
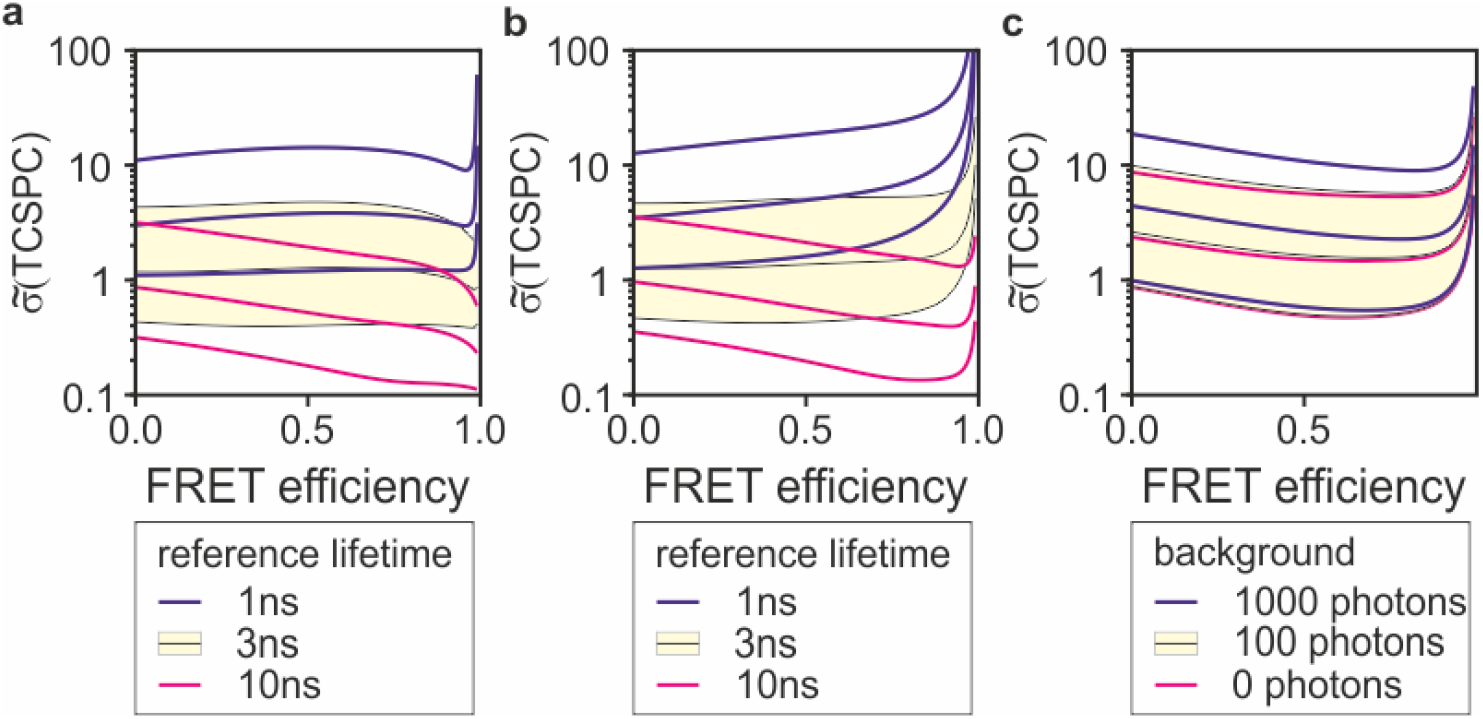
Photon-economy in FRET estimation by TCSPC. The statistical errors normalized to photon-budget for FRET measured by FLIM was estimated numerically. (a) Photon-efficiency of TCSPC for an ideal TCSPC system with Dirac-like IRF for an unquenched lifetime of 1ns (blue), 3ns (black lines and yellow area) and 10ns (magenta). Curves of the same colour represent the cases where the fractional contribution of the FRET-fraction is 10%, 50% and ~90% from top to bottom. (b) Same simulations as shown in panel (a) but with a finite IRF of 38ps full-width at half-maximum. (c) Simulations for a 3ns unquenched fluorescence lifetime, where an uncorrelated background signal of 0 (magenta), 100 (black curves and yellow area) and 1,000 photons (blue) are considered. In all simulations, the number of photons emitted from the unquenched component is kept to 1,000 but the results shown do not change with the absolute number of total photons.

All the intermediate cases that are highlighted with the yellow surface in fig. 1 are also used as reference in fig. 2–4. To guide in the interpretation of the figures, we provide a practical example with a donor fluorophore exhibiting an unquenched fluorescence lifetime of 3ns, 50% FRET efficiency and with only half of the photons emitted by donor interacting with acceptor molecules. Under these conditions 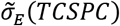 is approximately equal to one (fig. 1a). With a photon-budget equal to 1,000 photons, an ideal TCSPC would thus estimate a FRET efficiency of 50±3% (the standard deviation is 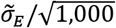). With a 90% of the photons emitted by the quenched donor, 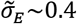 and the estimate of FRET efficiency would be around 50±1%. However, it should be noted that in the first case, around one quarter of photons are lost because of FRET, in the second case, almost half. Therefore, at even exposure times and not at even photon counts, the total photon counts in the second case would amount to ~750, resulting in an estimate of 50±4%. This example illustrates how the results shown are simple to interpret but, at the same time, the specific case of TCSPC presented in fig. 1 is compounded to loss of photons towards the acceptor fluorophore, photons which information is not used.

**Fig. 2.**
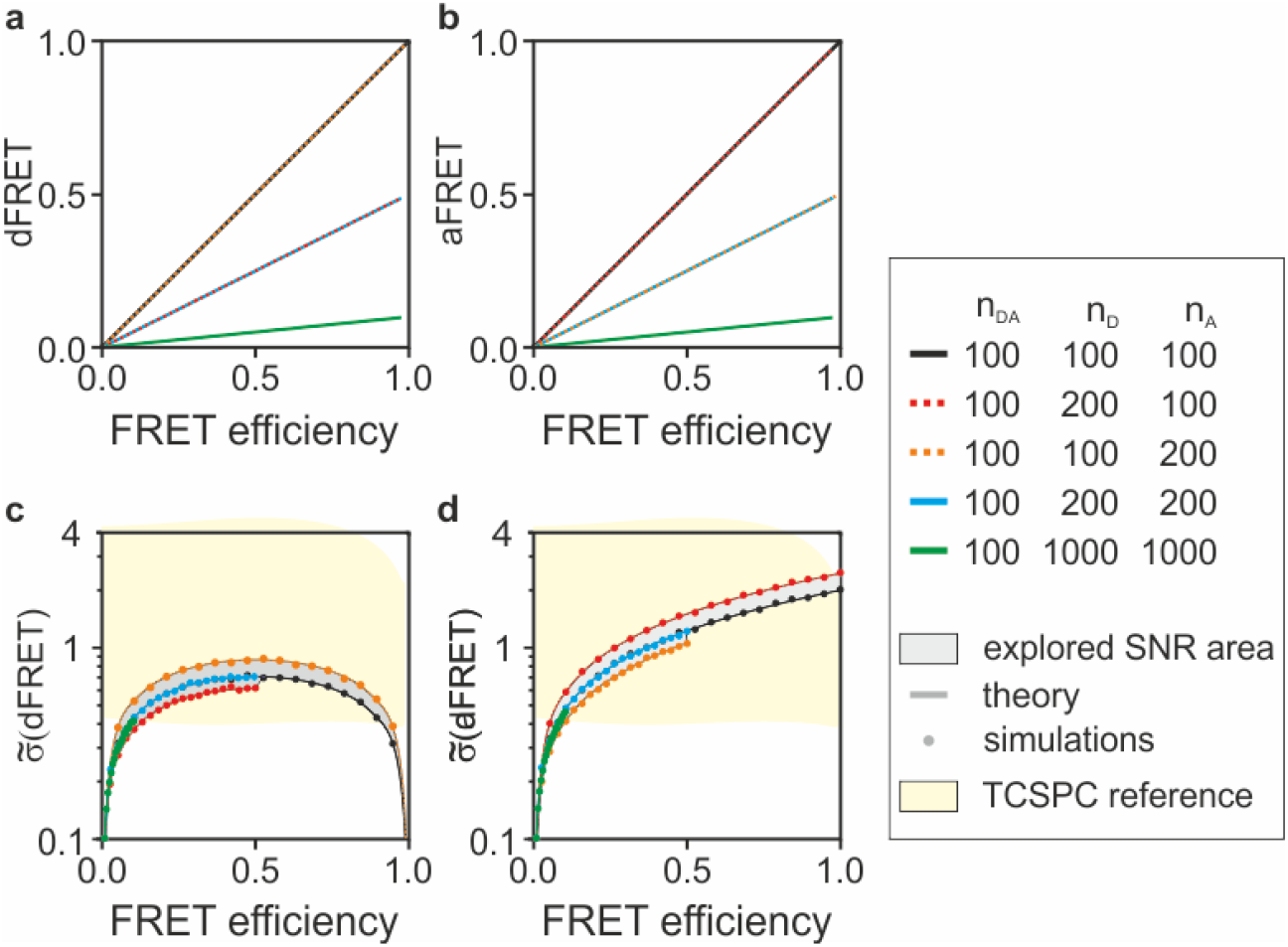
seFRET in the absence of background. The dFRET (a) and the aFRET (b) estimators are unbiased in the absence of background signals. When all molecules participate to FRET (black curves), both dFRET and aFRET measures FRET efficiency and trace the diagonal of the plots. dFRET and aFRET are insensitive to an excess of the non-interacting acceptor (orange) or donor (red) molecules, respectively. In all other conditions, for instance, partial participation of molecules to FRET (50% in blue and 10%in green), dFRET and aFRET measure lower FRET values but correct relative to their definitions (f_D_E and f_A_E, respectively). The normalized standard deviations for dFRET (c) and aFRET (d) vary with a sweep of the parameters (n_DA_, n_D_, n_A_ and E) albeit in a narrow SNR area (gray) and with a perfect match between the analytical solutions (dark gray curves) and the numerical simulations (solid circles). In yellow, the reference area explored by TCSPC from Fig. 1a is shown.

### C. Photon-economy of seFRET in the absence of cross-talks

First we consider the case where only intrinsic noise is present (Eqs. 10 and 13) with the proportionality constants η and ε set to 1, to aid the interpretation of the results. Fig. 2 shows numerical simulations (see Methods for details) carried out with one-hundred donor-acceptor pairs participating in Förster-type energy transfer (from 0 % to 100 % FRET efficiency) in the presence and absence of donor and acceptor molecules that do not undergo energy transfer (f_D_ = 10-100%, f_A_ = 10-100%,). The close match between the simulations and the analytical solutions demonstrates the consistency of the formalism we obtained. Fig. 2a-b shows that dFRET and aFRET are unbiased estimators for f_D_E and f_A_E, respectively. Furthermore, fig. 2c-d shows that the signal-to-noise ratio (SNR) in dFRET is always equal or better than aFRET. In these ideal conditions, dFRET is infinitely precise both in the absence of FRET or in the presence of 100% FRET efficiencies as the absence of signal from either the donor or acceptor channel unequivocally inform about the occurrence of these particular cases. The SNR values for dFRET and aFRET depend on the relative number of acceptors and donors in the sample; however, the estimators are quite robust in the absence of spurious signals. Indeed, seFRET explores a relatively narrow SNR area when varying the values of f_D_ and f_A_ (Fig. 2c-d - grey area). In comparison, the SNR for an ideal TCSPC (fig. 1a and fig. 2c-d, yellow area) explores a much wider area at the varying contribution of donors interacting with acceptor fluorophores.

### D. Photon-economy of seFRET in the presence of cross-talk

To illustrate the general principles underlying photon-statistics in seFRET and for an initial validation of the theoretical framework, in section C we have illustrated an ideal case that does not occur in realistic experimental conditions aiming. Next, we introduce spectral cross-talks (non-negligible AER and DER values) to evaluate at which extent these non-idealities degrade the efficiency of seFRET. Table 1 shows values that are reported in the literature for cross-talk parameters for a confocal (system 1) and a wide-field (system 2) microscope using typical yellow and cyan fluorescent proteins [22, 23].

Fig. 3a-b shows that dFRET and aFRET are unbiased estimators also in the presence of cross-talk when the calibration parameters are properly measured. However, fig. 3c-d shows that the noise performances of the estimators are significantly deteriorated in the presence of cross-talk, accounting for a twenty-fold (system 2, blue curve and circles) and a five-fold (system 1, red curves and circles) increase of standard deviations compared to the ideal measurement (black curves and circles are those also shown in fig. 2).

**Fig. 3.**
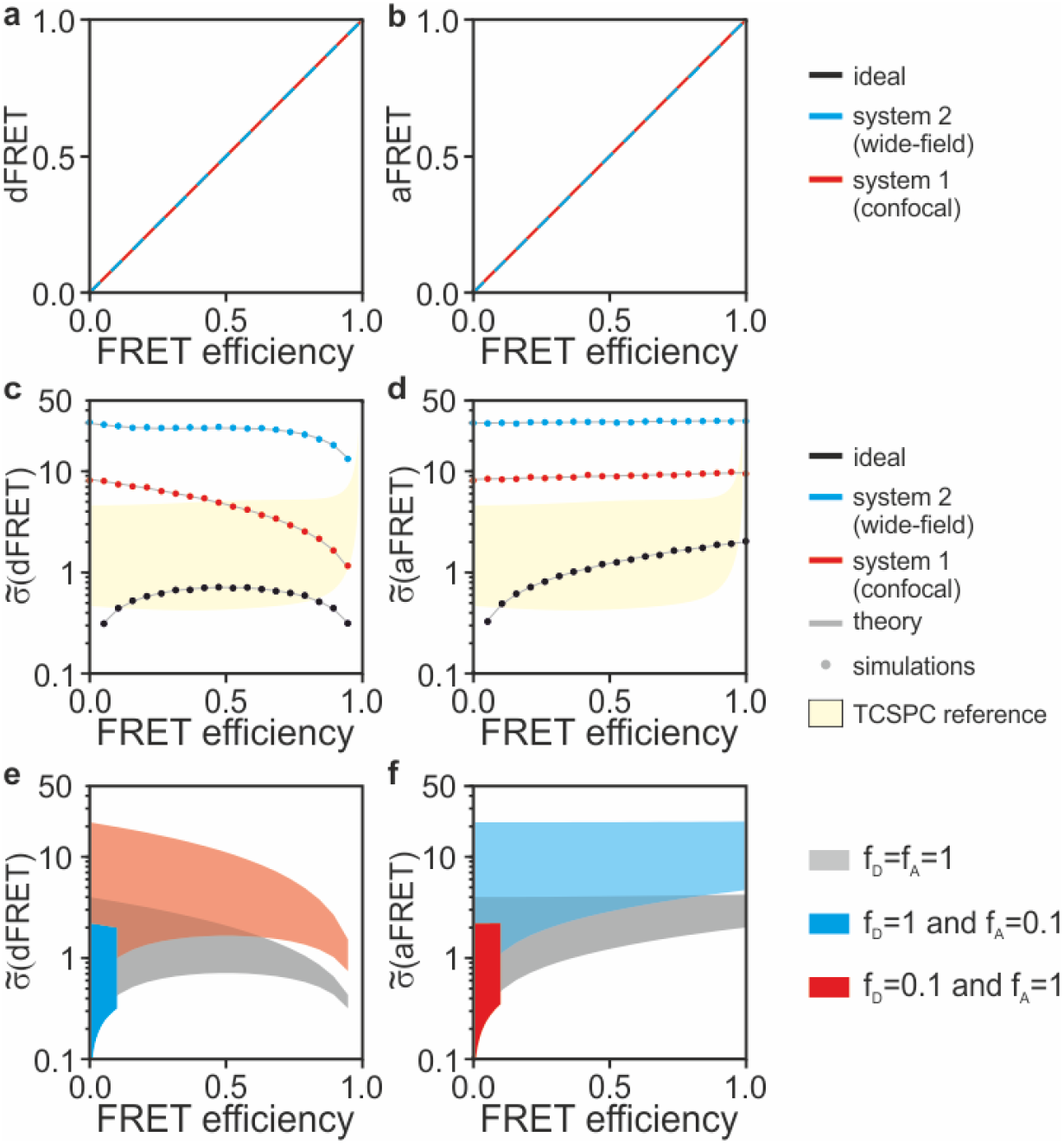
seFRET in the presence of spectral bleed through. Even in the presence of spectral cross-talk between channels, dFRET (a) and aFRET (b) are unbiased estimators as shown using the cross-talk reported in table one for a representative configuration of a confocal (system 1, red) and a wide-field (system 2, blue) microscope. However, cross-talk cause a significant deterioration of the SNR of the estimators (c-d). The loss of SNR and its dependency on DER, AER and the fraction of interacting donor/acceptor fluorophores is further illustrated in (e-f) where the SNR regions for molecules that all interact (grey), or where a minority of donor (red) or acceptor (blue) molecules interact is shown by varying DER and AER from 0 to 1. In yellow, the reference area explored by TCSPC from Fig. 1b is shown.

To generalize these results, in fig. 3e-f we show noise performance as a function of a parameter sweep, where we varied the AER and DER values from 0 to 1 with η and ε set to 1, similarly to fig. 1. We also simulated three conditions where: all molecules participate to FRET (f_D_=f_A_=1, grey area) and only a minority of donor (f_D_=0.1, f_A_=1, red area) or acceptor (f_D_=1, f_A_=0.1, blue area) molecules contribute to FRET. A decrease of interacting fractions causes significant deterioration of the SNR compared to the case where all molecules are interacting. Lower fractions of donor or acceptor molecules interacting causes the highest loss of efficiency to donor or acceptor estimators, respectively.

Donor imaging by FLIM does not suffer from spectral bleed-through and it is rather robust also to non-idealities such as broadening of the IRF (fig. 3 c-d, yellow areas). In realistic conditions, FRET estimates by TCSPC tend to outperform seFRET methods.

### E. seFRET in the presence of a background signal

We also studied how an unspecific background signal deteriorates the performances of dFRET and aFRET. Fig. 4 a-b shows that both dFRET and aFRET are biased and provide inaccurate estimations for FRET efficiency in the presence of background. For simplicity, the numerical simulations are carried out assuming an equal relative background contribution in all channels setting the number of background photons (B^DD^, B^AA^ and B^DA^) equal to a fixed fraction of photons emitted by donor and acceptor molecules (shown from 0% to 60%). In these conditions, dFRET overestimates FRET efficiencies at lower values (<50%) and underestimates energy transfer at higher values while aFRET overestimates FRET efficiencies with particularly large biases at low FRET values. These inaccuracies can be ameliorated by experimental corrections and, whenever possible, operating in conditions of high signal-to-background ratio. More importantly, the presence of background deteriorates the SNR of the dFRET and aFRET estimations. In the illustrative cases shown in fig. 4c-d, we set a background signal at 20% level showing the significant deterioration of SNR (note the logarithmic scale).

**Fig. 4.**
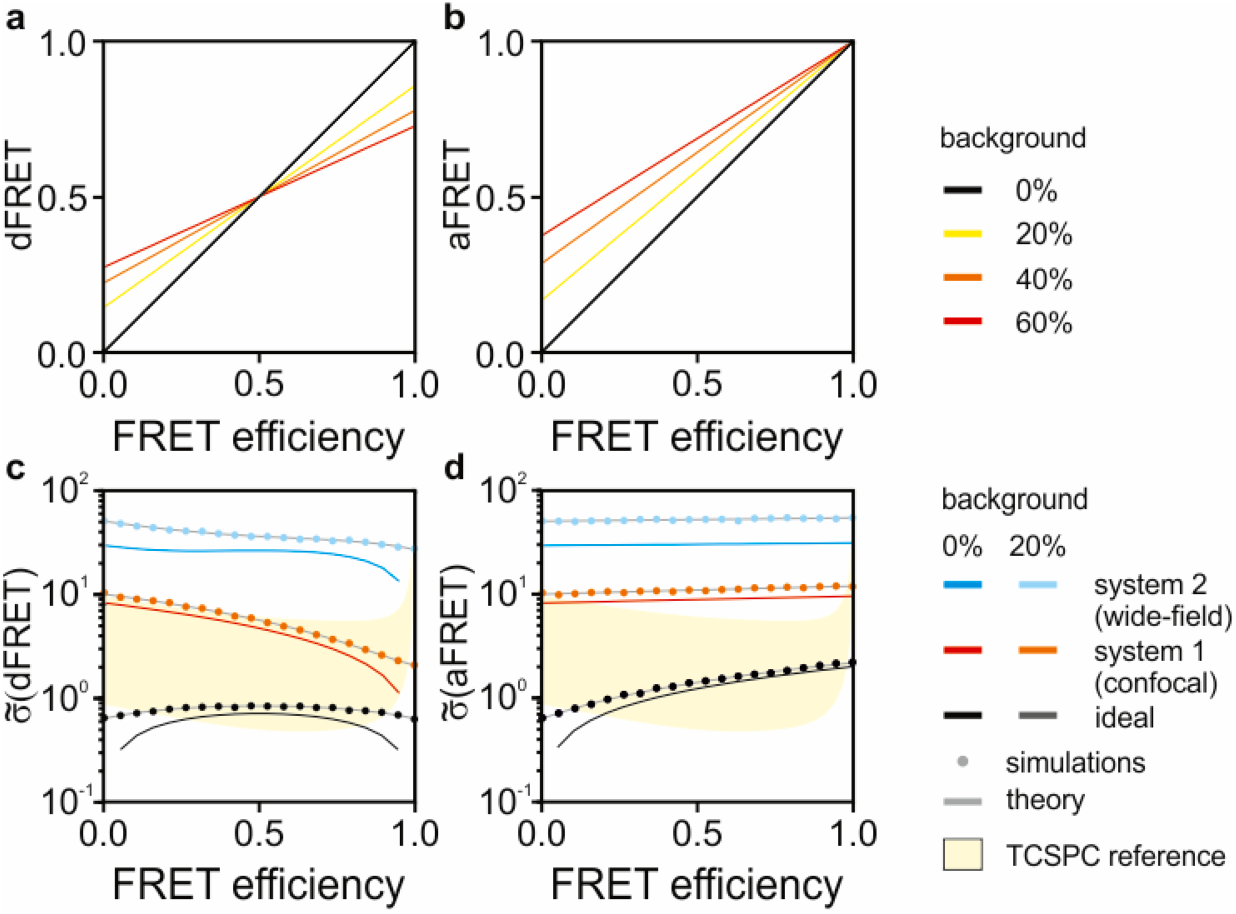
seFRET in the presence of background. When the measured fluorescence intensities are contaminated with an unspecific background signal, the dFRET (a) and aFRET (b) are not accurate estimators of f_D_E and f_A_E. The background-to-signal ratio simulated here (0%, black; 20%, yellow; 40%, orange and 60%, red) span a very broad range to illustrate the magnitude of the loss of accuracy. The analytical solutions (solid lines) describing the noise in dFRET (c) and aFRET (d) match the numerical simulations (solid circles) also in the presence of a background signal. We compare the noise for the systems also shown in fig. 1, *i.e.* system 1 (confocal, 0% background in red, 20% background in orange), system 2 (wide-field, 0% background in blue, 20% background in cyan) and the ideal case as a reference (0% background in black and 20% background in dark gray). In yellow, the reference area explored by TCSPC from Fig. 1c is shown.

Fluorescence signals from non-specific stains (*e.g.*, autofluorescence) deteriorate estimate obtained by any techniques. However, when the background is caused by instrumentation (*e.g.*, dark current or stray light), TCSPC is very robust to background noise, as illustrated by the yellow areas in fig. 3c-d. The lower and the upper boundaries of the yellow area corresponds to a signal to background ratio for TCSPC equal to infinity and to 0.5, respectively, while the signal to background noise simulated for seFRET is set to either infinite (no background) or 0.8.

## 3. Discussion

FRET imaging is a powerful method to probe cell biochemistry. Therefore, it is not surprising that a multitude of assays have been developed using FRET, from *in vitro* single-molecule detection [20] to *in vivo* imaging [34, 35], including both very common applications (*e.g.*, quantitative polymerase chain reaction and related hybridization-based assays [36]) and more specialist uses such as the study of protein conformations, interactions and post-translational modifications [4]. Because of its versatility, a plethora of methodologies for FRET imaging and data analysis [37] exists. Fluorescence lifetime imaging microscopy and sensitized emission FRET are two of the most common quantitative methods used for biochemical imaging. The choice between FLIM and seFRET often depends on the availability of specialist instrumentation (*e.g.*, for TCSPC/FLIM) or requirements such as fast acquisition speed (typically better for seFRET). FLIM in general - and TCSPC among FLIM techniques - is regarded as the most robust technique for FRET estimation [9]. In fact, FLIM requires fewer control samples and provides robust and reproducible absolute measurements across independent experimental sessions. The photon-statistics of the various implementations of fluorescence lifetime imaging have been studied in-depth (for example, in [10, 11, 13, 38–41]). Photon-counting techniques such as TCSPC are very efficient and can provide the highest attainable SNR. Taken together, these observations suggest that TCSPC/FLIM is the most accurate and precise technique to quantify FRET, a conclusion also supported by some experimental observations [9].

Therefore, seFRET is commonly used as a fast, simpler and more cost-effective alternative. Breakthroughs in FLIM-enabling technologies [42–48] and data analysis [49–51] are reducing the barrier to adoption for FLIM; therefore, as the choice between the two techniques might be slowly drift away from technical requirements, we aimed to develop a comparative analysis of the limits of both techniques from an information theory perspectives to provide guidance on the selection and also optimization of these methodologies.

Interestingly, seFRET can outperform TCSPC in the estimation of FRET in the ideal conditions of negligible spectral cross-talk. In this specific case, TCSPC can attain higher SNR only when the majority (>50%) of donor fluorophores are engaged in FRET with acceptors. In all other cases, the dFRET estimator performs significantly better. A better photon-efficiency of the dFRET estimator for FRET efficiency results from the capability of dFRET to utilize information from photons emitted from both donor and acceptor fluorophores. However, the higher precision of dFRET is vastly reduced as soon as realistic levels of spectral cross-talks and background are taken into account. Furthermore, we did not consider the additional statistical and systematic errors that the reference measurements required by seFRET causes and other sources of noise manifesting in detectors that do not operate in single-photon counting. Therefore, despite the excellent performance of seFRET compared to TCSPC, TCSPC might generally outperform seFRET in reproducibility, accuracy and precision in practical implementations. However, it is important to note that the appropriate optimization of imaging parameters for seFRET can make seFRET rather competitive also for its high precision, something that might be often underestimated. Moreover, the use of long-Stokes shift of acceptor fluorophores for seFRET, not usually implemented to the best of our knowledge, might result in vast improvements in the SNR of this intensity-based technique. We also note that we compared seFRET to TCSPC as an established gold-standard in FRET detection. Other FLIM implementations can provide significantly worse performances than TCSPC [17, 38] and TCSPC can as well deteriorate its performance when pushed to competitive fast detection limits [52, 53], suggesting that there are cases when FLIM can lose its competitive edge relative to a simpler seFRET technique.

Ultimately, one of the most substantial differences between FLIM and seFRET is that FLIM is typically used for the detection of donor fluorescence, permitting researchers to streamline the use of the visible spectrum or to optimize Foster distances with dark acceptors [54, 55], avoiding cross-talks and issues related to chromatic aberrations. On the contrary, seFRET uses the complete photon-budget emitted by the donor and acceptor fluorophores providing a mean to improve precision. Therefore, in those cases where the benefits of a dark chromophore might be irrelevant, the combination of seFRET and TCSPC (*e.g.*, in dual-colour FLIM) might provide significant improvements in precision in FRET estimation. A higher precision leads directly to an improvement in the capability to resolve smaller biochemical differences in living cells. From a theoretical standpoint, this improvement in biochemical resolving power can also be understood for the general analysis of Fisher information in multi-dimensional or multi-parametric detection systems (see for example the photon partitioning theorem in [56, 57]). From a practical point of view, dual-colour fast high-resolution FLIM might be increasingly accessible thanks to the ongoing revolution in time-resolved detection technologies and could provide yet unexplored ideal performances.

## 4. Methods

All the analytical solutions were obtained manually, but their consistency was evaluated with the use of Mathematica (Wolfram). The numerical simulations and plots were generated with bespoke software written in Matlab (Mathworks) and available at the GitHub repository github.com/alesposito/FisherInformation with a CC BY 4.0 license. The Cramer-Rao lower bound for TCSPC was obtained with parameters sweeps utilizing the computational core developed Bouchet and colleagues [32]. Briefly, we utilized their methods to compute the standard deviation, normalized to the total (donor) photon counts, of the shorter fluorescence lifetime estimate. This was the fluorescence lifetime quenched via FRET and evaluated as a double-exponential fit with constant background and known IRF. Fig. 1a was generated considering 1,000 photons emitted by donor molecules not participating in FRET with a reference lifetime equal to 1, 3 or 10ns. Both the reference lifetime and the number of photons were used as fixed parameters. The number of photons emitted by quenched donors was varied from 100, 500 to 10,000 corresponding to SNRs of 0.1, 0.5 and 10. Energy transfer efficiency was varied from 0 to 100% in 128 steps on a power series. We used TCSPC as a gold-standard reference and, therefore, we utilized parameters of high-end systems. The laser repetition rate was set to 80MHz with a histogram resolution of 8-bits resulting in a bin time resolution of 48.8ps. Fig. 1b was generated in the same way, but using an experimental IRF provided by Bouchet et al. [32], with a nominal IRF of 38ps full-width at half-maximum. For Fig. 1c, we simulated only a reference fluorescence lifetime of 3ns. All other parameters the same as in Fig. 1b, we varied the number of photons in an uncorrelated background for Fig. 1c, including 0, 100 and 1,000 photons, which had to be estimated. This parameter sweep resulted in different signal-to-background ratios as described in the result sections. The results are shown as normalized by the total photon count emitted, that is including the sum of photons emitted by the FRET-fraction, the unquenched fraction and the background.

Numerical simulations for seFRET were utilized to propagate errors in the unmixing equations described in this work. First, we synthesized noiseless images using the same mathematical framework; subsequently, we added Poissonian noise and unmixed the images to determine how noise propagates to the FRET estimates aFRET and dFRET. The results for both TCSPC and seFRET depend on the ratio of the emitted photons and not on the absolute values (not shown) of photon counts, thus they are presented as normalized to the total photon counts for generality. All the simulations are computationally light and compatible with older versions of Matlab and computers; however, the results reported in this work were performed with Dell Precision workstation equipped with an Intel Xeon CPU E5-1620v3 and 64GB of RAM and Matlab 2018a.

## 5. Acknowledgements

We acknowledge funding from the Medical Research Council core grants (MC_UU_12022/1 and MC_UU_12022/8) to Prof. Ashok Venkitaraman (MRC Cancer Unit at the University of Cambridge, UK).

