## Supporting Information for "Photon-statistics in sensitized emission FRET and FLIM: a comparative theoretical analysis"

Alessandro Esposito[1,*]

[1]MRC Cancer Unit, University of Cambridge, Biomedical Campus, Cambridge, CB20XY, UK

*

### Supporting information

**1. Fisher information** | A brief introduction to Fisher information theory

**2. Fisher information matrix and seFRET** | Applications of Fisher theory to seFRET

**3. Comparison between different nomenclatures** | A guide to differences between nomenclatures

#### 1. Fisher information

The Fisher information matrix defines the information content of parameter estimators accordingly to stochastic models of the experiments. The Fisher information matrix for fluorescence lifetime sensing has been derived analytically and studied previously [15-17]. The Fisher information matrix for sensitized emission FRET was described, to our knowledge, only for the case of single-molecule detection [22]. The information content of an experiment can be described by the likelihood function (L):

$$L = \prod_{i=1}^{m} \frac{[F(iU)-F((i-1)U)]^{N_i} e^{-[F(iU)-F((i-1)U)]}}{N_i!} \quad \textbf{(S1)}$$

Here, we broadly adhere to the formalism described by Hall and colleagues [16] revised with some nomenclature introduced by Watkins and Yang [22]. The likelihood function is the product of the probability to detect $N_i$ (i=1, …, m) photons in $m$ independent channels, photons that are Poisson distributed with a rate defined by the model function F:

$$F(\vec{u},\vec{q}) = \int_0^{\vec{u}} f(\vec{v},q)d\vec{v} \quad \textbf{(S2)}$$

where $f(u,q)$ is the expected average signal as a function of channel parameters $u$ and experimental parameters that should be estimated $q$. For instance, for time-resolved detection, $u$ will equal $t$ and integration will be carried out over time bins. $U$ is the bandwidth of the channel (*e.g.*, spectral bandwidth, time-gate width). $f$ can be factorized into parameters such as the excitation rate ($k_{ex}$), the integration time of the signal $T$, and two functions that depend on the experimental parameters ($\zeta(q)$) and on the detection system ($S(u)$):

$$f(\vec{u},\vec{q}) = k_{ex}TS(\vec{u})\zeta(\vec{q}) \quad \textbf{(S3)}$$

With distributions commonly found in spectroscopy, the elements $j_{ij}$ of the Fisher information matrix can be derived computing the negative of the expectation of each second derivative of $f$:

$$j_{ij} = -E\left[\frac{\partial^2 \ln f(\vec{u},\vec{q})}{\partial q_i \partial q_j}\right]_{\vec{u}} \quad \textbf{(S4)}$$

Watkins and Yang [22] showed that when the signal is described by the Poisson statistics, the equation above further simplifies to:

$$j_{jk} = T\sum_{i=1}^{m} \frac{k_i S_i(\vec{u})}{\zeta_i(\vec{q})} \frac{\partial \zeta_i(\vec{q})}{\partial q_j} \frac{\partial \zeta_i(\vec{q})}{\partial q_k} \quad \textbf{(S5)}$$

In the next sections, we will describe the algebraic manipulations of seFRET formalism required to obtain the separation of variables shown in Eq. S3 that will permit the estimation of Eq. S5. The Fisher information matrix is essential when studying the noise performance of an estimator as the the Cramer-Rao theorem states that the lower bounds of the variance of the unbiased estimators of $q$ are defined by the inverse of the Fisher information matrix.

#### 2. Fisher information matrix and seFRET

The fluorescence emission as a function of wavelenght (λ) for the case of seFRET can be described as the sum of photons emitted by the donor fluorophore, photons emitted by sensitised acceptors (SE) and photons emitted by acceptors upon direct excitation (DE) with the donor excitation light source:

$$f(\lambda, E, n_D, n_A, n_{DA}) = k_{ex}T\{S_D(\lambda)[n_D + n_{DA}(1-E)] + n_{DA}S_{SE}(\lambda)E + n_A S_{DE}(\lambda)\} \quad \textbf{(S6)}$$

$S_D$, $S_{SE}$ and $S_{DE}$ include spectral characteristics of the fluorophores and detection system, *e.g.*, quantum yields, molar extinction coefficients, spectral overlaps; $n_D$, $n_A$ and $n_{DA}$ are the number of non-interacting donor molecules, non-interacting acceptor molecules and interacting donor-acceptor pairs in the sample, respectively. seFRET detected by the typical three-filter method is carried out with two acquisition channels for donor emission and sensitized emission that will effectively carry information on energy transfer and a reference acquisition channel with direct excitation of the acceptor. The main difference between single molecule seFRET and seFRET imaging is the requirement in the latter to reference the measured FRET signal to the concentration of the acceptor molecules. In order to compute the Fisher information matrix, we shall integrate eq. S6:

$$F(\lambda) = \int_0^\lambda \{k_{ex}T\{S_D(\lambda)[n_D + n_{DA}(1-E)] + n_{DA}S_{SE}(\lambda)E + n_A S_{DE}(\lambda)\}\}d\lambda \quad \textbf{(S7)}$$

$$F(\lambda) = k_{ex}T\left\{[n_D + n_{DA}(1-E)]\int_0^\lambda S_D(\lambda)d\lambda + n_{DA}E\int_0^\lambda S_{SE}(\lambda)d\lambda + n_A\int_0^\lambda S_{DE}(\lambda)d\lambda\right\} \quad \textbf{(S8)}$$

Using Eq. S8, we can estimate the fluorescence intensity collected within a detection channel optimized for donor emission fluorescence of spectral window [$\lambda_{d1}$, $\lambda_{d2}$]:

$$I^{DD} = F(\lambda_{d2}) - F(\lambda_{d1}) \quad \textbf{(S9)}$$

$$I^{DD} = k_{ex}T\left\{[n_D + n_{DA}(1-E)]\int_{\lambda_{d1}}^{\lambda_{d2}} S_D(\lambda)d\lambda + n_{DA}E\int_{\lambda_{d1}}^{\lambda_{d2}} S_{SE}(\lambda)d\lambda + n_A\int_{\lambda_{d1}}^{\lambda_{d2}} S_{DE}(\lambda)d\lambda\right\} \quad \textbf{(S10)}$$

$$I^{DD} = k_{ex}TS_{DD}[n_D + n_{DA}(1-E)] \quad \textbf{(S11)}$$

The cross-talk between the donor channel and acceptor emission is assumed to be negligible:

$$\begin{cases} \int_{\lambda_{d1}}^{\lambda_{d2}} S_D(\lambda)d\lambda = S_{DD} \\ \int_{\lambda_{d1}}^{\lambda_{d2}} S_{SE}(\lambda)d\lambda = 0 \\ \int_{\lambda_{d1}}^{\lambda_{d2}} S_{DE}(\lambda)d\lambda = 0 \end{cases} \quad \textbf{(S12)}$$

It is convenient also to model an unspecific background signal and to introduce $f_A$ and $f_D$, the fractions of interacting donor or acceptors:

$$I^{DD} = k_{ex}T\{S_{DD}[n_D + n_{DA}(1-E)] + B_{DD}\} = k_{ex}T\{S_{DD}N_D[1 - f_D E] + B_{DD}\} \quad \textbf{(S13)}$$

Where $N_D$ is the total number of donor fluorophores. In a similar way, the intensity collected in the donor channel can be described as a function of $f_A$ and the total number of acceptor fluorophores ($N_A$). In summary:

$$I^{DD}(f_D E) = k_{ex}T[S_{DD}N_D(1 - f_D E) + B_{DD}] \quad \textbf{(S14)}$$

$$I^{DD}(f_A E) = k_{ex}T[S_{DD}N_D - S_{DD}N_A f_A E + B_{DD}] \quad \textbf{(S15)}$$

Similarly, we can evaluate the intensity collected in the sensitized emission channel by integration of eq. S8 over the spectral range [$\lambda_{a1}$, $\lambda_{a2}$]:

$$I^{DA} = F(\lambda_{a2}) - F(\lambda_{a1}) \quad \textbf{(S16)}$$

$$I^{DA} = k_{ex}T\left\{[n_D + n_{DA}(1-E)]\int_{\lambda_{a1}}^{\lambda_{a2}} S_D(\lambda)d\lambda + n_{DA}E\int_{\lambda_{a1}}^{\lambda_{a2}} S_{SE}(\lambda)d\lambda + (n_A + n_{DA})\int_{\lambda_{a1}}^{\lambda_{a2}} S_{DE}(\lambda)d\lambda\right\} \quad \textbf{(S17)}$$

Using eq. S18:

$$\begin{cases} \int_{\lambda_{a1}}^{\lambda_{a2}} S_D(\lambda)d\lambda = S_{DD}DER \\ \int_{\lambda_{a1}}^{\lambda_{a2}} S_{SE}(\lambda)d\lambda = S_{AA} \\ \int_{\lambda_{a1}}^{\lambda_{a2}} S_{DE}(\lambda)d\lambda = \varepsilon S_{AA}AER \end{cases} \quad \textbf{(S18)}$$

ε is a proportionality factor between the intensity of the excitation light used for donor and acceptor excitation. DER = $[I^{DA}/I^{DD}]_{only\text{-}donor}$ is the donor emission ratio, a control measurement for the donor spectral bleed-through into the acceptor channel using a donor-only control sample performed by measuring the proportion of signal in the acceptor channel relative to the donor channel, and estimating the ratio between the intensities detected in the acceptor. AER = $[I^{DA}/I^{AA}]_{only\text{-}acceptor}$ is the acceptor excitation ratio, a control measurement aimed to estimate the direct excitation of acceptor fluorophores by measuring the proportion of intensities in the acceptor channel with a donor-only sample with excitation light optimal for donor and acceptor excitation, respectively. Adding an unspecific channel background $B_{DA}$, we can write:

$$I^{DA} = F(\lambda_{a2}) - F(\lambda_{a1}) \quad \textbf{(S19)}$$

$$I^{DA} = k_{ex}T\{[n_D + n_{DA}(1-E)]S_{DD}DER + [n_{DA}E + (n_A + n_{DA})\varepsilon AER]S_{AA} + B_{DA}\} \quad \textbf{(S20)}$$

Which can be rewritten using the definitions of $f_D$, $N_D$ and $N_A$ to:

$$I^{DA} = k_{ex}T\{n_{DA}(S_{AA} - S_{DD}DER)E + \varepsilon AER(n_A + n_{DA})S_{AA} + S_{DD}DER(n_D + n_{DA}) + B_{DA}\} \quad \textbf{(S21)}$$

$$I^{DA} = k_{ex}T\{N_D(S_{AA} - S_{DD}DER)f_DE + S_{AA}N_A\varepsilon AER + S_{DD}N_DDER + B_{DA}\} \quad \textbf{(S22)}$$

We can apply an analogous parametrization with $f_A$ to obtain:

$$I^{DA}(f_DE) = k_{ex}T\{N_D(S_{AA} - S_{DD}DER)f_DE + S_{AA}N_A\varepsilon AER + S_{DD}N_DDER + B_{DA}\} \quad \textbf{(S23)}$$

$$I^{DA}(f_AE) = k_{ex}T\{N_A(S_{AA} - S_{DD}DER)f_AE + S_{AA}N_A\varepsilon AER + S_{DD}N_DDER + B_{DA}\} \quad \textbf{(S24)}$$

The intensity value measured in the acceptor reference channel ($I^{AA}$) does not depend on energy transfer or the number of donor molecules and can be simply described as:

$$I^{AA} = k_{ex}T\varepsilon(S_{AA}N_A + B_{AA}) \quad \textbf{(S25)}$$

We can simplify the description of the Fisher information matrix using the following set of substitutions:

$$\begin{cases} E_D = f_DE \\ E_A = f_AE \\ C_D = S_{DD}N_D \\ C_A = S_{AA}N_A \\ \eta = {S_{DD}}/{S_{AA}} \\ N_P = k_{ex}T \end{cases} \quad \textbf{(S26)}$$

$E_D$ and $E_A$ are the apparent FRET efficiencies measured in seFRET by the donor (dFRET) and acceptor (aFRET) normalized estimators; $C_D$ and $C_A$ are the relative concentrations of donor and acceptor in arbitrary units; $\eta$ is the ratio of the relative brightness of the donor and the acceptor fluorophores. Substituting the set of definition shown in eq. S14-15 and S23-25, we thus obtain:

$$
\begin{cases}
I^{DD}(E_D, C_D) = N_P[C_D(1-E_D) + B_{DD}] \\
I^{DD}(E_A, C_D, C_A) = N_P[C_D - \eta C_A E_A + B_{DD}] \\
I^{DA}(E_D, C_D, C_A) = N_P[(\eta^{-1} - DER)C_D E_D + C_A \varepsilon AER + C_D DER + B_{DA}] \\
I^{DA}(E_A, C_D, C_A) = N_P[(1 - DER\eta)C_A E_A + C_A \varepsilon AER + C_D DER + B_{DA}] \\
I^{AA}(C_A) = N_P \varepsilon [C_A + B_{AA}]
\end{cases}
\tag{S27}
$$

Following the formalism introduced by Watkins and colleagues, we can rewrite Eq. S27 with the following parametrization:

$$
\begin{cases}
I^{DD}(E_D, C_D, C_A) = F^{DD}[(1-\beta_{DD}^{-1})\zeta_{DD}(E_D, C_D, C_A) + \beta_{DD}^{-1}] \\
F^{DD} = N_P(1 + B_{DD}) \\
\beta_{DD} = \frac{1+B_{DD}}{B_{DD}} \\
\zeta_{DD}(E_D, C_D, C_A) = C_D(1-E_D)
\end{cases}
\tag{S28}
$$

$$
\begin{cases}
I^{DD}(E_A, C_D, C_A) = F^{DD}[(1-\beta_{DD}^{-1})\zeta_{DD}(E_A, C_D, C_A) + \beta_{DD}^{-1}] \\
F^{DD} = N_P(1 + B_{DD}) \\
\beta_{DD} = \frac{1+B_{DD}}{B_{DD}} \\
\zeta_{DD}(E_A, C_D, C_A) = C_D - \eta C_A E_A
\end{cases}
\tag{S29}
$$

$$
\begin{cases}
I^{DA}(E_D, C_D, C_A) = F^{DA}[(1-\beta_{DA}^{-1})\zeta_{DA}(E_D, C_D, C_A) + \beta_{DA}^{-1}] \\
F^{DA} = N_P(1 + B_{DA}) \\
\beta_{DA} = \frac{1+B_{DA}}{B_{DA}} \\
\zeta_{DA}(E_D, C_D, C_A) = (\eta^{-1} - DER)C_D E_D + C_A \varepsilon AER + C_D DER
\end{cases}
\tag{S30}
$$

$$
\begin{cases}
I^{DA}(E_A, C_D, C_A) = F^{DA}[(1-\beta_{DA}^{-1})\zeta_{DA}(E_A, C_D, C_A) + \beta_{DA}^{-1}] \\
F^{DA} = N_P(1 + B_{DA}) \\
\beta_{DA} = \frac{1+B_{DA}}{B_{DA}} \\
\zeta_{DA}(E_A, C_D, C_A) = (1 - DER\eta)C_A E_A + C_A \varepsilon AER + C_D DER
\end{cases}
\tag{S31}
$$

$$
\begin{cases}
I^{AA}(C_A) = F^{AA}[(1-\beta_{AA}^{-1})\zeta_{AA}(C_A) + \beta_{AA}^{-1}] \\
F^{AA} = \varepsilon N_P(1 + B_{AA}) \\
\beta_{AA} = \frac{1+B_{AA}}{B_{AA}} \\
\zeta_{AA}(C_A) = C_A
\end{cases}
\tag{S32}
$$

The Fisher information matrix can be now estimated by computing a set of derivatives of the functions $\zeta$s and substituting in Eq. S5:

$$
\begin{pmatrix}
\frac{\partial \zeta_{DD}}{\partial E_D} & \frac{\partial \zeta_{DD}}{\partial C_D} & \frac{\partial \zeta_{DD}}{\partial C_A} \\
\frac{\partial \zeta_{DA}}{\partial E_D} & \frac{\partial \zeta_{DA}}{\partial C_D} & \frac{\partial \zeta_{DA}}{\partial C_A} \\
\frac{\partial \zeta_{AA}}{\partial E_D} & \frac{\partial \zeta_{AA}}{\partial C_D} & \frac{\partial \zeta_{AA}}{\partial C_A}
\end{pmatrix}
=
\begin{pmatrix}
-C_D & 1 & 0 \\
(\eta^{-1} - DER)C_D & DER - (\eta^{-1} - DER)E_D & \varepsilon AER \\
0 & 0 & 1
\end{pmatrix}
\tag{S33}
$$

$$\begin{pmatrix} \frac{\partial \zeta_{DD}}{\partial E_A} & \frac{\partial \zeta_{DD}}{\partial C_D} & \frac{\partial \zeta_{DD}}{\partial C_A} \\ \frac{\partial \zeta_{DA}}{\partial E_A} & \frac{\partial \zeta_{DA}}{\partial C_D} & \frac{\partial \zeta_{DA}}{\partial C_A} \\ \frac{\partial \zeta_{AA}}{\partial E_A} & \frac{\partial \zeta_{AA}}{\partial C_D} & \frac{\partial \zeta_{AA}}{\partial C_A} \end{pmatrix} = \begin{pmatrix} -\eta C_A & 1 & -\eta E_A \\ (1 - DER\eta)C_A & DER & \varepsilon AER + (1 - DER\eta)E_A \\ 0 & 0 & 1 \end{pmatrix} \quad \textbf{(S34)}$$

Each element of the Fisher information matrix can be then computed accordingly to Eq. S5:

$$J_{11} = N_P C_D^2 \left[\frac{1}{C_D(1-E_D)+B_{DD}} + \frac{\eta^{-1}-DER}{(\eta^{-1}-DER)C_D E_D + C_A \varepsilon AER + C_D DER + B_{DA}}\right] \quad \textbf{(S35)}$$

$$J_{12} = N_P C_D \left\{-\frac{1}{C_D(1-E_D)+B_{DD}} + \frac{(\eta^{-1}-DER)C_D[DER-(\eta^{-1}-DER)E_D]}{(\eta^{-1}-DER)C_D E_D + C_A \varepsilon AER + C_D DER + B_{DA}}\right\} \quad \textbf{(S36)}$$

$$J_{13} = N_P C_D \frac{(\eta^{-1}-DER)\varepsilon AER}{(\eta^{-1}-DER)C_D E_D + C_A \varepsilon AER + C_D DER + B_{DA}} \quad \textbf{(S37)}$$

$$J_{21} = J_{12} \quad \textbf{(S38)}$$

$$J_{22} = N_P \left\{\frac{1}{C_D(1-E_D)+B_{DD}} + \frac{[DER-(\eta^{-1}-DER)E_D]^2}{(\eta^{-1}-DER)C_D E_D + C_A \varepsilon AER + C_D DER + B_{DA}}\right\} \quad \textbf{(S39)}$$

$$J_{23} = N_P \left\{\frac{[DER-(\eta^{-1}-DER)E_D]\varepsilon AER}{(\eta^{-1}-DER)C_D E_D + C_A \varepsilon AER + C_D DER + B_{DA}}\right\} \quad \textbf{(S40)}$$

$$J_{31} = J_{13} \quad \textbf{(S41)}$$

$$J_{32} = J_{23} \quad \textbf{(S42)}$$

$$J_{33} = N_P \left\{\frac{1}{C_A + B_{AA}}\right\} \quad \textbf{(S43)}$$

Here, we show explicitly only the evaluation of the Fisher information matrix related to dFRET but similar algebraic steps can be used also for aFRET. The Cramer-Rao bound for the variance of $E_D$ is the first element of the inverse matrix with elements described in Eq. S35-S43.

$$\begin{aligned} \sigma_E^2(dFRET) = N_P^{-1}\{ & B_{DD}[DER\,\eta\,(1-E_D) + E_D]^2 C_D^{-2} + \\ & B_{DA}[\eta\,(1-E_D)]^2 C_D^{-2} + \\ & B_{AA}[\,AER\,\eta\,(1-E_D)]^2 C_D^{-2}\varepsilon + \\ & AER\,(AER+1)[C_A(1-E_D)^2\eta^2]C_D^{-2}\varepsilon + \\ & DER[(DER+1)(1-E_D)\eta + 2E_D][(1-E_D)^2\eta]C_D^{-1} + \\ & E_D(1-E_D)[E_D(1-\eta)+\eta]C_D^{-1}\} \end{aligned} \quad \textbf{(S44)}$$

The variance in the FRET efficiency estimate is equal to the sum of a background variance ($\sigma_B^2$), a variance which depends on the spectral bleed through ($\sigma_{SBT}^2$) and a variance which does not depends on background contributions ($\sigma_E^2$):

$$\sigma_{E_D}^2 = \frac{\tilde{\sigma}_B^2 + \tilde{\sigma}_{SBT}^2 + \tilde{\sigma}_E^2}{N_P} \quad \textbf{(S45)}$$

The tilde indicates variances normalized to the Poisson noise variance ($N_P$). Finally, we can write simpler analytical solutions for each component:

$$\begin{cases} \tilde{\sigma}_E^2(dFRET) = \tilde{\sigma}_{B_{DD}}^2(dFRET) + \tilde{\sigma}_{B_{DA}}^2(dFRET) + \tilde{\sigma}_{B_{AA}}^2(dFRET) \\ \tilde{\sigma}_{B_{DD}}^2(dFRET) = B_{DD}[DER\,\eta\,(1-E_D) + E_D]^2 C_D^{-2} \\ \tilde{\sigma}_{B_{DA}}^2(dFRET) = B_{DA}[\eta\,(1-E_D)]^2 C_D^{-2} \\ \tilde{\sigma}_{B_{AA}}^2(dFRET) = B_{AA}[AER\,\eta\,(1-E_D)]^2 C_D^{-2}\varepsilon \end{cases} \quad \textbf{(S46)}$$

$$\begin{cases} \tilde{\sigma}_{SBT}^2(dFRET) = \tilde{\sigma}_{DER}^2(dFRET) + \tilde{\sigma}_{AER}^2(dFRET) \\ \tilde{\sigma}_{DER}^2(dFRET) = AER(AER+1)[C_A(1-E_D)^2\eta^2]C_D^{-2}\varepsilon \\ \tilde{\sigma}_{AER}^2(dFRET) = DER[(DER+1)(1-E_D)\eta + 2E_D](1-E_D)^2\eta C_D^{-1} \end{cases} \quad \textbf{(S47)}$$

$$\tilde{\sigma}_E^2(dFRET) = E_D(1-E_D)[E_D(1-\eta)+\eta]C_D^{-1} \quad \textbf{(S48)}$$

Similarly, we can evaluate the analytical descriptions for the noise of the estimator aFRET:

$$\begin{cases} \tilde{\sigma}_B^2(aFRET) = \tilde{\sigma}_{B_{DD}}^2(aFRET) + \tilde{\sigma}_{B_{DA}}^2(aFRET) + \tilde{\sigma}_{B_{AA}}^2(aFRET) \\ \tilde{\sigma}_{B_{DD}}^2(aFRET) = B_{DD}\mathrm{DER}\, C_A^{-2} \\ \tilde{\sigma}_{B_{DA}}^2(aFRET) = B_{DA}C_A^{-2} \\ \tilde{\sigma}_{B_{AA}}^2(aFRET) = B_{AA}[\varepsilon\, AER\, E_A]^2 C_D^{-2}\varepsilon^{-1} \end{cases} \quad \textbf{(S49)}$$

$$\begin{cases} \tilde{\sigma}_{SBT}^2(aFRET) = \tilde{\sigma}_{DER}^2(aFRET) + \tilde{\sigma}_{AER}^2(aFRET) \\ \tilde{\sigma}_{DER}^2(aFRET) = DER(DER+1)[C_D - C_A E_A \eta]C_A^{-2} \\ \tilde{\sigma}_{AER}^2(aFRET) = AER[(AER+1)\varepsilon + 2E_A]C_A^{-1} \end{cases} \quad \textbf{(S50)}$$

$$\tilde{\sigma}_E^2(aFRET) = E_A(1+\varepsilon^{-1}\mathrm{A})C_A^{-1} \quad \textbf{(S51)}$$

#### 3. Comparison between different nomenclatures

| | **AER** | **DER** | **α** | **β** |
|---|---|---|---|---|
| In this work | AER | DER | ηDER | $(AER\varepsilon)^{-1}$ |
| In Hoppe *et al.* | α | β | $\xi\beta\gamma^{-1}$ | $\gamma^{-1}$ |

**Table S1.** Conversion of nomenclature from Elder *et al.*

| | **AER** | **DER** | **η** | **ε** |
|---|---|---|---|---|
| In Elder *et al.* | AER | DER | α/DER | β/AER |
| In Hoppe *et al.* | α | β | $\xi\gamma^{-1}$ | $(\alpha\gamma)^{-1}$ |

**Table S2**. Conversion of nomenclature from this work

| | **α** | **β** | **ξ** | **γ** |
|---|---|---|---|---|
| In this work | AER | DER | $\eta(AER\varepsilon)^{-1}$ | AERε |
| In Elder *et al.* | AER | DER | $\alpha(DER\beta)^{-1}$ | $\beta^{-1}$ |

**Table S3.** Conversion of nomenclature from Hoppe *et al.*
